# Predicting Capsid Protein Binding Sites in Single-Stranded RNA Viruses Using Machine Learning from Local Geometric Features

**DOI:** 10.64898/2026.09.10.750674

**Authors:** Yunong Melody Wu, Junjie Zhang

**Affiliations:** Rye Country Day School, Rye, NY 10580; Center for Phage Technology, Department of Biochemistry and Biophysics, Texas A&M University, College Station, TX 77843

## Abstract

Selective recognition of viral RNA by capsid proteins is essential for genome packaging during the assembly of single-stranded RNA (ssRNA) viruses. However, identification of capsid protein binding sites in the RNA genome remains challenging because current experimental techniques are labor-intensive and low-throughput, motivating the development of computational approaches. Here, we present a sequence-based framework that integrates RNA tertiary structural modeling, local geometric feature extraction, and machine learning to predict capsid protein binding sites. Using the Qβ bacteriophage as a proof-of-concept system, we constructed a benchmark dataset of experimentally identified binding and non-binding RNA fragments. We designed a set of geometric descriptors to characterize the local structural features of the RNA backbone. When repeatedly trained and tested with these geometric descriptors on different subsets of the benchmark dataset, the neural network showed a strong ability to distinguish capsid protein binding sites from non-binding RNA fragments, achieving an area under the receiver operating characteristic curve (AUC) of 0.88 in a 5-fold cross-validation. To evaluate whether the model can predict RNA binding sites without experimental structures, we applied it to local geometric features derived from AlphaFold-predicted RNA structures. Despite substantial structural differences between predicted and experimentally determined models, the classifier retained considerable predictive performance (AUC = 0.75), indicating that approximate RNA tertiary structures may still preserve biologically meaningful information for capsid binding site prediction. Furthermore, the failed predictions suggest that viral genome packaging is not only governed by intrinsic RNA structural features, but also by additional dynamic factors beyond static RNA conformations. In summary, our findings provide new mechanistic insights into RNA–capsid interactions and establish a foundation for extending this approach to diverse ssRNA viruses.

## 1. Introduction

Single-stranded RNA (ssRNA) viruses constitute one of the largest groups of viruses, infecting organisms across all domains of life [1]. They are responsible for numerous infectious diseases with substantial impacts on global health, including COVID-19 [2], influenza [3], and hepatitis C [4]. Additionally, ssRNA viruses have become valuable platforms for the development of RNA-based therapeutics and vaccines [5]. During viral assembly, the viral RNA genome is enclosed within a protective protein shell known as the capsid [6]. Viral genome packaging is a highly regulated process that requires specific interactions between viral capsid proteins and selected regions of the RNA genome. Understanding the molecular principles governing these RNA–capsid interactions is fundamental to elucidating viral assembly mechanisms and enabling the rational engineering of antiviral strategies and RNA delivery systems.

These RNA–capsid interactions are mediated by specific regions within the viral RNA genome, known as packaging signals, which are recognized by viral capsid proteins to ensure efficient viral assembly [7]. In this study, these regions are referred to as capsid protein binding sites. Experimentally, capsid protein binding sites can be identified using approaches such as cryo-electron microscopy, X-ray crystallography, crosslinking-based sequencing, and biochemical assays [8]. However, these methods are labor-intensive, low-throughput, and have been applied to only a limited number of viruses. Computational approaches have therefore emerged as attractive alternatives, ranging from sequence motif analysis [9] to structure-based modeling [10]. However, sequence-based methods often overlook the three-dimensional organization of RNA. On the other hand, structure-based approaches typically require accurate structural information, which remains unavailable or difficult to obtain for most viruses. Consequently, there remains a need for computational methods that accurately predict capsid protein binding sites by integrating both sequence and structural information directly from viral RNA sequences.

Recent advances in artificial intelligence have substantially improved RNA structure prediction from sequence [11]. Although current prediction methods do not yet achieve experimental-level accuracy, they often preserve local geometric characteristics and topological relationships that are biologically meaningful. We hypothesized that these approximate structural representations contain sufficient information to identify capsid protein binding sites with the assistance of machine learning. Based on this hypothesis, we developed a computational framework that predicts capsid protein binding sites directly from viral RNA sequences by first generating RNA tertiary structures computationally, extracting local geometric features from the predicted structures to be evaluated in a supervised neural network model.

Using the Qβ bacteriophage [23] as a proof-of-concept system, we demonstrate the feasibility of this strategy for predicting capsid protein binding sites. Our results show that local geometric features derived from predicted RNA structures provide measurable predictive power for identifying capsid protein binding sites, indicating that approximate RNA tertiary structures preserve biologically meaningful signals for capsid recognition. At the same time, the incomplete predictive performance suggests that selective viral genome packaging is governed not only by static RNA structure but also by RNA conformational dynamics and the molecular context of viral assembly. Overall, this framework offers a new strategy for integrating sequence and structural information to investigate RNA–capsid interactions and lays the foundation for extending this approach to other single-stranded RNA viruses.

## 2. Model and Methods

### 2.1. Overview of the computational framework

We developed and tested a computational framework for predicting capsid protein binding sites in single-stranded RNA viruses by integrating local geometric features and supervised machine learning (**Figure 1a**). The framework consists of four major steps. First, experimentally determined RNA–capsid interactions from the Qβ bacteriophage were used to construct a benchmark dataset of positive and negative RNA fragments. Second, local geometric descriptors were extracted from each RNA fragment using packing angles and dihedral angles calculated from RNA backbone phosphate atoms. Third, these geometric features were used to train a feedforward neural network classifier to distinguish capsid protein binding sites from non-binding regions. Finally, we evaluated the model on the Qβ sequence alone, without relying on an experimentally determined structure. Local geometric features were extracted directly from its computationally predicted structure, and the trained neural network was used to predict capsid protein binding sites across the RNA genome.

**Figure 1.**
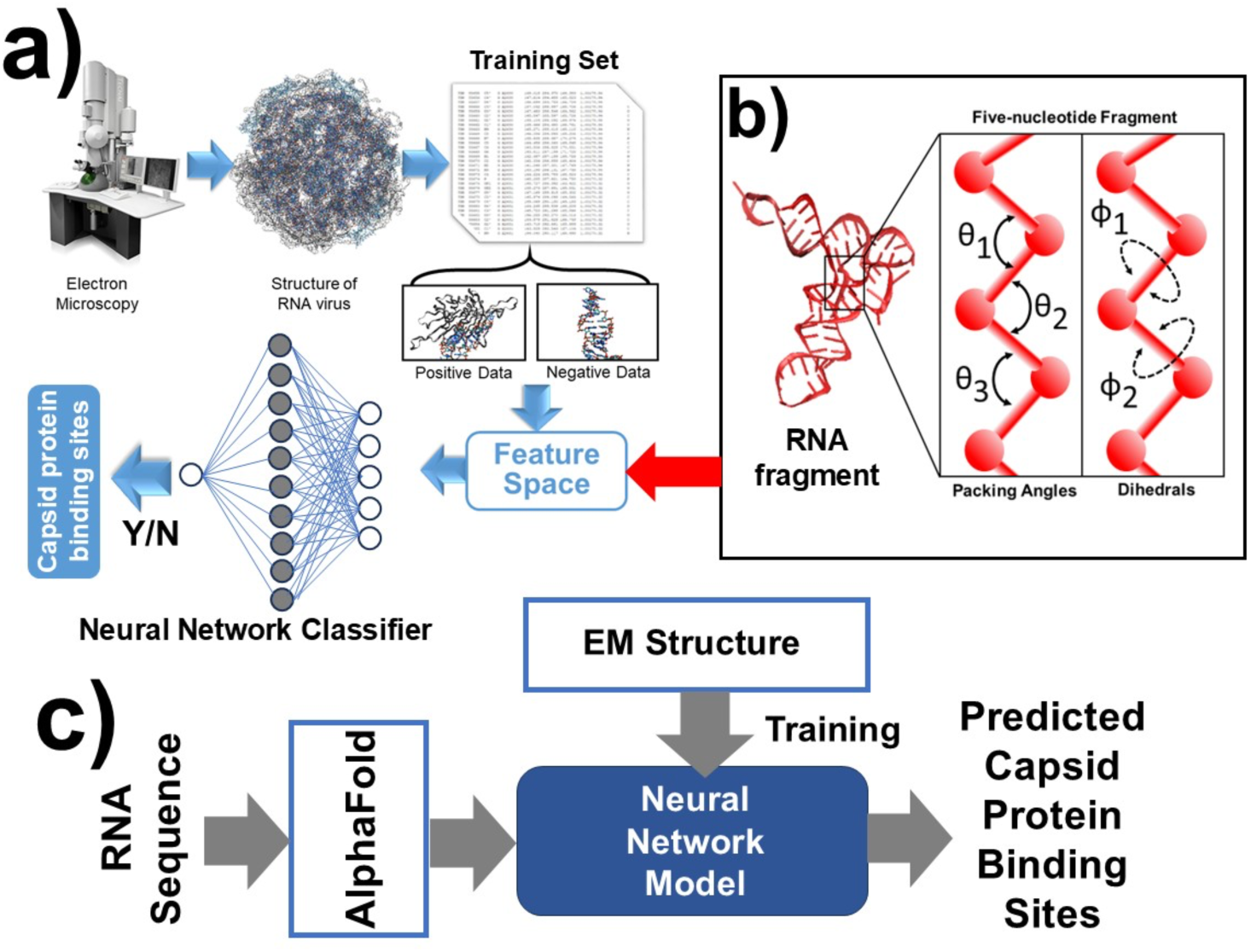
Overview of the computational framework. **(a)** Workflow of the machine learning pipeline. The RNA structure of the Qβ bacteriophage is used as a training set. Local geometric features were extracted from the RNA fragments and fed into a supervised neural network. Ground-truth binding labels were obtained from experimentally determined RNA–capsid contacts in the cryo-EM structure. **(b)** Illustration of the local geometric features extracted from a five-nucleotide RNA fragment. Three packing angles (θ₁–θ₃) and two dihedral angles (φ₁–φ₂) were calculated from consecutive phosphate atoms to characterize the local RNA geometry. **(c)** The neural network was trained using experimentally annotated binding and non-binding nucleotides. After training, we did a blind test. Pretending we don’t know the experimentally derived structure, we submitted the RNA sequence of Qβ to the AlphaFold server. The geometric features were then derived from the structural model. They were input to the trained neural network to predict the probability of capsid protein binding for each nucleotide in the RNA genome.

### 2.2. Benchmark dataset construction

The benchmark dataset was constructed using the experimentally determined structure of the single-stranded RNA bacteriophage Qβ, recently solved by cryo-electron microscopy [12]. The viral genome consists of 4,127 nucleotides enclosed within a near-icosahedral capsid. To identify capsid protein binding sites, every nucleotide in the viral genome was examined for physical contacts with surrounding capsid proteins. For simplicity, the minimum Euclidean distance between every atom in the nucleotide and every atom in the capsid proteins was calculated. A nucleotide was considered to interact with a capsid protein if the minimum atom–atom distance was smaller than a predefined distance cutoff (5.0 Å). Nucleotides satisfying this criterion were labeled as capsid protein binding sites, whereas all remaining nucleotides were considered non-binding positions.

Using this procedure, 100 experimentally supported capsid protein binding sites were identified throughout the Qβ RNA genome. To construct a balanced benchmark dataset for supervised learning, an equal number of RNA fragments that did not interact with capsid proteins were randomly selected as negative samples. Each sample consisted of a five-nucleotide RNA fragment centered on the corresponding nucleotide, resulting in a benchmark dataset containing 100 positive and 100 negative samples.

### 2.3. Local geometric feature construction

To characterize the local three-dimensional geometry of each RNA fragment, we designed a five-dimensional geometric feature representation based on the RNA backbone (**Figure 1b**). Each RNA fragment consisted of five consecutive nucleotides, and the positions of their phosphate (P) atoms were used to calculate geometric descriptors. Three packing angles were calculated from every three consecutive phosphate atoms within the fragment, describing the local bending of the RNA backbone. In addition, two dihedral angles were calculated from every four consecutive phosphate atoms, capturing the local torsional geometry of the RNA chain. Consequently, each five-nucleotide fragment was represented by a five-dimensional feature vector,

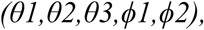

where *θ* denotes packing angles and *ϕ* denotes dihedral angles. These geometric descriptors provide a compact representation of local RNA conformation while remaining independent of the global orientation of the RNA molecule.

### 2.4. Neural network classifier

A supervised feedforward neural network [13] was implemented to classify RNA fragments as capsid protein binding sites or non-binding sites based on the extracted geometric features. The network consisted of an input layer containing five neurons corresponding to the five geometric features. The input layer was connected to a fully connected hidden layer containing ten neurons with rectified linear unit (ReLU) activation. The hidden layer was followed by a single output neuron with a sigmoid activation function, which generated a probability between 0 and 1, representing the likelihood that the input RNA fragment corresponds to a capsid protein binding site.

The network was trained as a binary classifier using the benchmark dataset described above. Positive samples corresponded to experimentally identified capsid protein binding sites, whereas negative samples represented RNA fragments without detectable protein contacts. Binary cross-entropy was used as the loss function, and network parameters were optimized using the Adam optimizer [14]. To evaluate the generalization performance of the model, we performed 5-fold cross-validation. The benchmark dataset was randomly divided into five approximately equal subsets. In each fold, four subsets (80% of the data) were used to train the neural network, while the remaining subset (20%) was held out for testing. This procedure was repeated five times, with each subset serving once as the test set, ensuring that every sample was evaluated by a model that was not trained on that sample. The prediction results from all five folds were then combined to evaluate the overall performance of the classifier. During training, the network parameters were iteratively updated to minimize the binary cross-entropy loss until convergence. The neural network was implemented using TensorFlow [15].

### 2.5. Prediction of capsid protein binding sites from RNA sequences without experimental data

To predict capsid protein binding sites for an RNA sequence alone (**Figure 1c**), the RNA tertiary structure was first predicted computationally using AlphaFold [16]. While we use the RNA genome sequence of Qβ, a notable divergence exists between the predicted and experimental tertiary structures (**Figure 3a**). This structural difference mimics realistic conditions where prediction errors occur, allowing us to rigorously evaluate the resilience of our machine-learning approach. This predicted RNA structure was then processed using the same geometric feature extraction procedure described above to calculate the three packing angles and two dihedral angles for every five-nucleotide fragment along the RNA sequence.

Each resulting five-dimensional feature vector was subsequently provided as input to the trained neural network, which generated the probability that the corresponding fragment represented a capsid protein binding site. By scanning the RNA sequence with a sliding five-nucleotide window, the framework produced nucleotide-level predictions of capsid protein binding sites across the entire viral genome.

### 2.6. Code and data availability

All relevant source codes can be found in the GitHub repository: https://github.com/melody144/CPBSpred.

## 3. Results

To investigate whether local RNA geometry contains information associated with capsid protein binding, we first compared the geometric features of RNA fragments corresponding to capsid protein binding sites with those of non-binding fragments. **Figure 2a** presents the distributions of the packing angles calculated from the positive and negative datasets. Although the two distributions partially overlap, their median values and interquartile ranges differ, indicating that RNA fragments participating in capsid protein interactions exhibit distinct local geometric characteristics. This observation suggests that capsid protein binding sites possess intrinsic structural signatures, e.g. mostly stem-loops, that can potentially be exploited for computational prediction.

**Figure 2.**
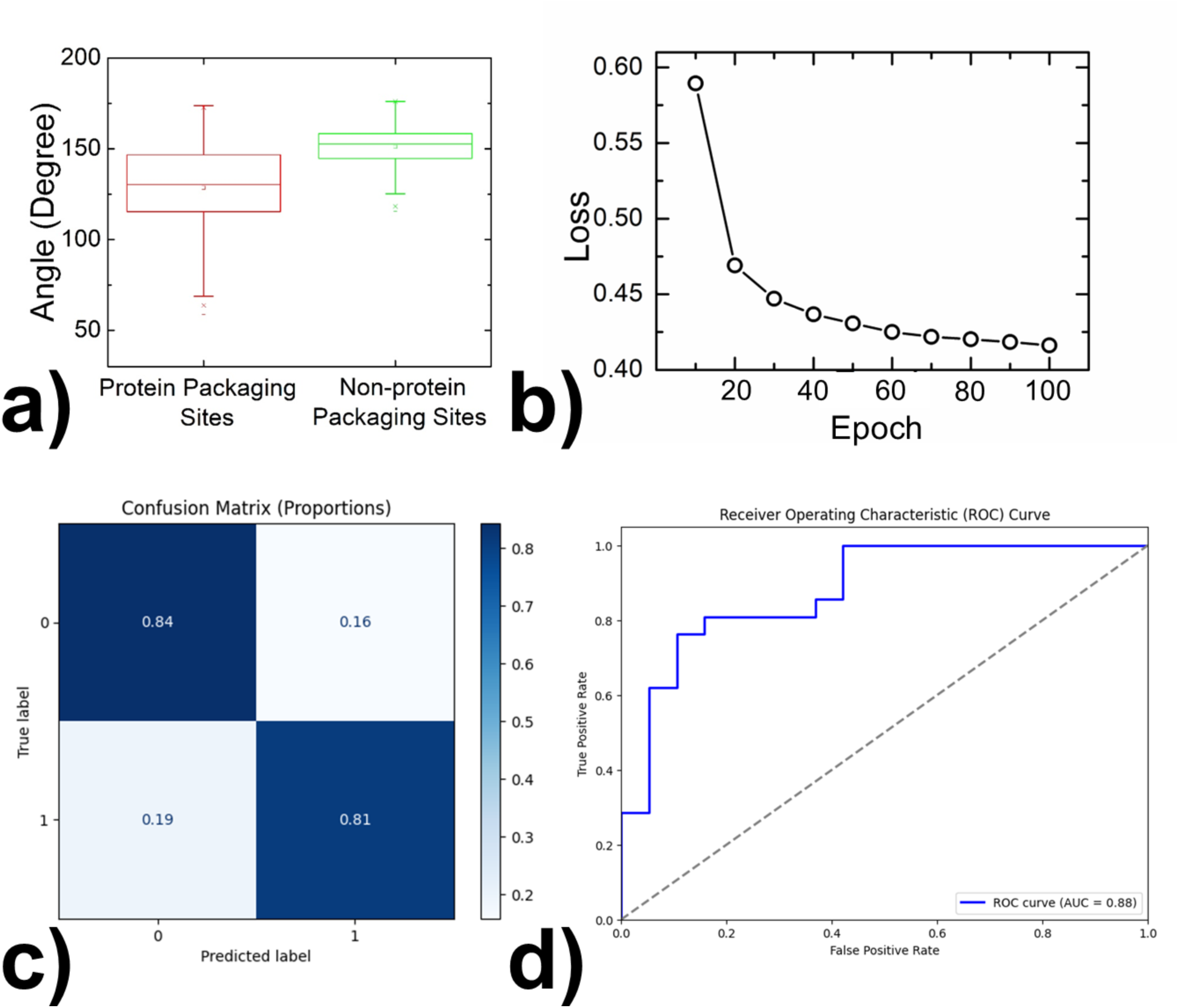
Performance evaluation of the neural network classifier. **(a)** Distribution of the local packing angles for RNA fragments corresponding to experimentally identified capsid protein binding sites (positive dataset) and non-binding sites (negative dataset). Boxes represent the interquartile range (25th–75th percentiles), horizontal lines indicate the median, whiskers extend to 1.5 times the interquartile range, and circles denote outliers. **(b)** Training loss of the neural network during optimization. **(c)** Confusion matrix obtained from 10-fold cross-validation. **(d)** Receiver operating characteristic (ROC) curve of the classifier.

Motivated by these observations, we trained a feedforward neural network using the five local geometric features extracted from each RNA fragment. Model performance was evaluated using 5-fold cross-validation. During training, the binary cross-entropy loss decreased rapidly over the first several training epochs and gradually converged to a stable value after approximately 40 epochs (**Figure 2b**). The smooth decrease in the loss function without noticeable oscillation demonstrates that the neural network successfully learned discriminative geometric features from the training data and achieved stable optimization without obvious signs of training instability.

The predictive performance of the trained classifier was first evaluated using the confusion matrix obtained from cross-validation (**Figure 2c**). The classifier correctly identified 84% of non-binding RNA fragments and 81% of experimentally determined capsid protein binding sites, while only 16% of non-binding fragments were incorrectly classified as binding sites and 19% of true binding sites were missed. These results indicate that the neural network achieved balanced classification performance for both positive and negative samples, demonstrating that the extracted local geometric features effectively distinguish capsid protein binding sites from other regions of the viral RNA genome.

We further evaluated the overall discrimination ability of the classifier using the receiver operating characteristic (ROC) curve [17] (**Figure 2d**). The ROC curve shows a model’s classification quality by graphing the True Positive Rate (TPR) against the False Positive Rate (FPR) across every possible decision threshold, where the TPR means the fraction of binding sites correctly recognized by our model among all the real binding sites, and the FPR means the fraction of non-binding sites that were wrongly classified as binding sites among all the non-binding sites [24]. The figure shows that our ROC curve lies substantially above the diagonal corresponding to random classification, indicating that the model consistently distinguishes binding sites from non-binding sites across different classification thresholds. The area under the ROC curve (AUC) reached 0.88, demonstrating excellent discriminative ability.

Collectively, these cross-validation results show that local geometric features extracted from RNA local backbone structures provide sufficient information for accurately identifying capsid protein binding sites and support the feasibility of using machine learning to capture the structural determinants of RNA–capsid interactions.

Having established that local geometric features derived from experimentally determined RNA structures can reasonably distinguish capsid protein binding sites, we next investigated whether the same framework could be applied when only the viral RNA sequence is available. To this end, we predicted the tertiary structure of the Qβ RNA genome using AlphaFold and compared the resulting model with the experimentally determined cryo-EM structure (**Figure 3a**). Although the overall architectures differ substantially, reflecting the current limitations of RNA tertiary structure prediction, the predicted structure provides a complete three-dimensional representation that can be analyzed computationally.

**Figure 3.**
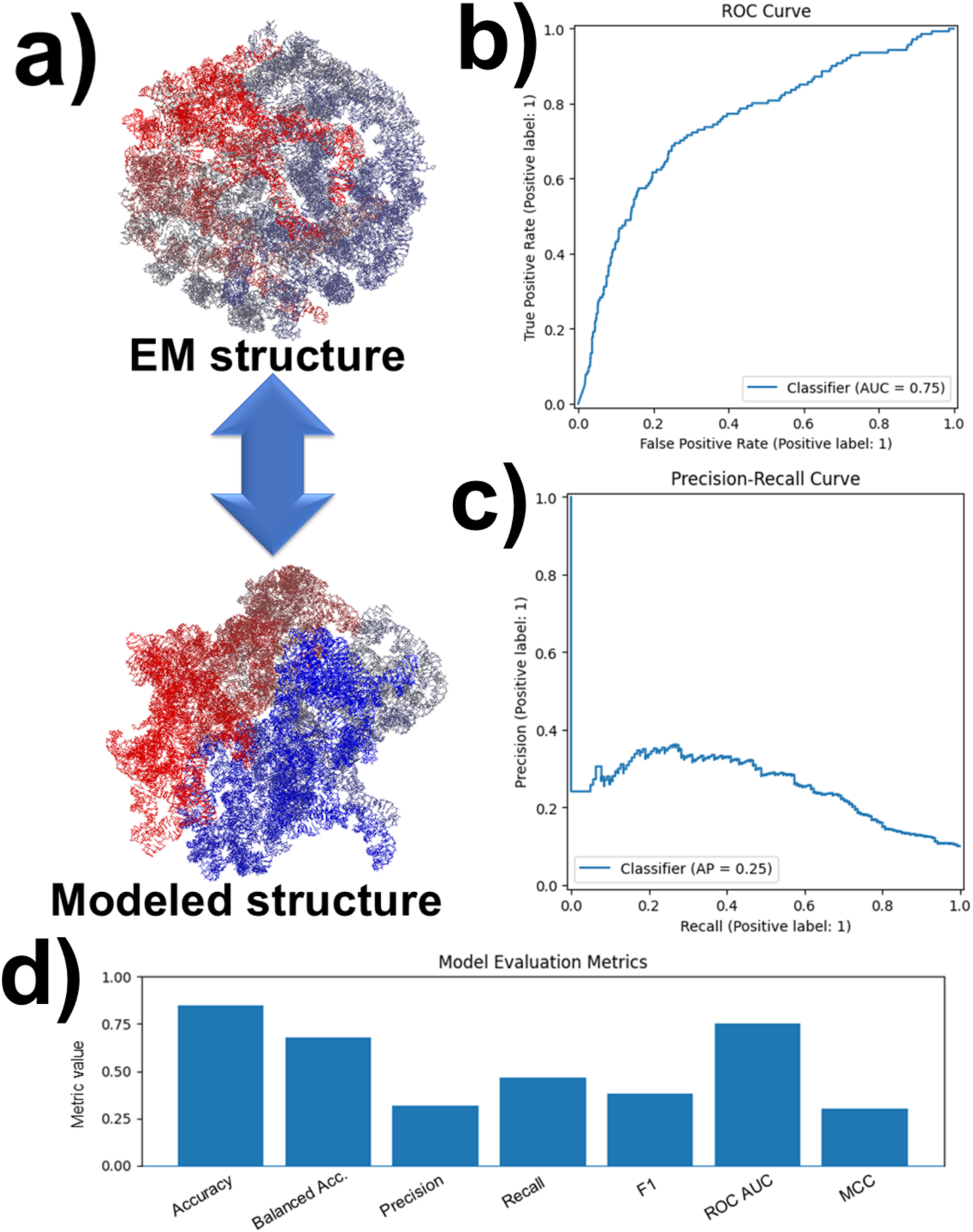
Prediction of capsid protein binding sites using AlphaFold-predicted RNA structures. **(a)** Comparison between the experimentally determined cryo-EM structure and the AlphaFold-predicted tertiary structure of the Qβ RNA genome. **(b)** Receiver operating characteristic (ROC) curve of capsid protein binding site prediction using geometric features extracted from the AlphaFold-predicted structure. **(c)** Precision–recall (PR) curve of the prediction model. **(d)** Performance metrics of the neural network classifier, including accuracy, balanced accuracy, precision, recall, F1 score, Matthews correlation coefficient (MCC), ROC AUC, and average precision (AP).

We next extracted the same five local geometric features from the AlphaFold-predicted RNA structure and applied the previously trained neural network to predict capsid protein binding sites across the Qβ genome. The predicted binding probabilities were then compared with experimentally determined binding sites identified from the cryo-EM structure. Despite the structural differences between the AlphaFold-predicted and experimental models, the neural network retained measurable predictive performance. The ROC curve yielded an area under the curve (AUC) of 0.75 (**Figure 3b**), indicating that the model can distinguish binding sites from non-binding sites substantially better than random classification. Consistent with this observation, the precision–recall curve exhibited an average precision (AP) of 0.25 (**Figure 3c**), demonstrating that the AlphaFold-predicted RNA structures preserve useful information for identifying capsid protein binding sites despite their limited structural accuracy. Additional evaluation metrics, including an overall accuracy of approximately 85%, balanced accuracy of 68%, F1 score [18] of 0.38, and Matthews correlation coefficient [19] of 0.30, further support the moderate predictive performance of the proposed framework (**Figure 3d**).

The intermediate prediction accuracy obtained using AlphaFold-derived RNA structures provides important biological insights into viral genome packaging. On one hand, the observed predictive signal indicates that computationally predicted RNA structures preserve functionally relevant information, including local geometric characteristics and folding patterns that contribute to capsid protein recognition. On the other hand, the reduced performance relative to experimentally determined structures suggests that static RNA tertiary structures alone do not fully determine binding specificity. Instead, selective viral genome packaging is likely governed by both intrinsic RNA structural features and additional biological factors that are not captured by current computational models. Capsid proteins may recognize transient RNA conformations, folding intermediates, or structural ensembles that are absent from a single predicted structure. Furthermore, viral assembly occurs within a dynamic molecular environment where RNA modifications, molecular chaperones, host or viral cofactors, ionic conditions, and genome compaction can all influence RNA–protein interactions.

Together, these findings support a hybrid model in which RNA tertiary structure provides an important, but incomplete, encoding of the viral packaging code. Future computational frameworks will likely require integrating static structural representations with RNA conformational dynamics, assembly kinetics, and cellular context to achieve a more comprehensive understanding of selective viral genome packaging.

To further examine the basis of individual predictions, we compared representative RNA fragments in the experimentally determined Qβ structure with their corresponding conformations in the AlphaFold-predicted model (**Figure 4**). In the two true-positive examples shown in **Figure 4a**, the experimentally identified capsid protein binding sites remained geometrically exposed in the predicted structure, allowing the neural network to correctly classify them as binding sites. In contrast, **Figure 4b** shows a false-negative example in which an experimentally confirmed binding fragment was positioned in the interior of the AlphaFold model. Because its predicted local geometry did not resemble an accessible capsid-contacting region, the model failed to identify it as a binding site. This discrepancy may arise because viral genome packaging is a dynamic assembly process in which RNA molecules undergo continuous conformational rearrangements during capsid formation. The RNA fragment may adopt an exposed conformation during assembly that facilitates capsid recognition but subsequently assumes a different conformation in the predicted static structure. On the other hand, **Figure 4c** illustrates a false-positive case: the RNA fragment does not contact the capsid in the cryo-EM structure but is located on the surface of the predicted structure, leading the classifier to predict it as a binding site.

**Figure 4.**
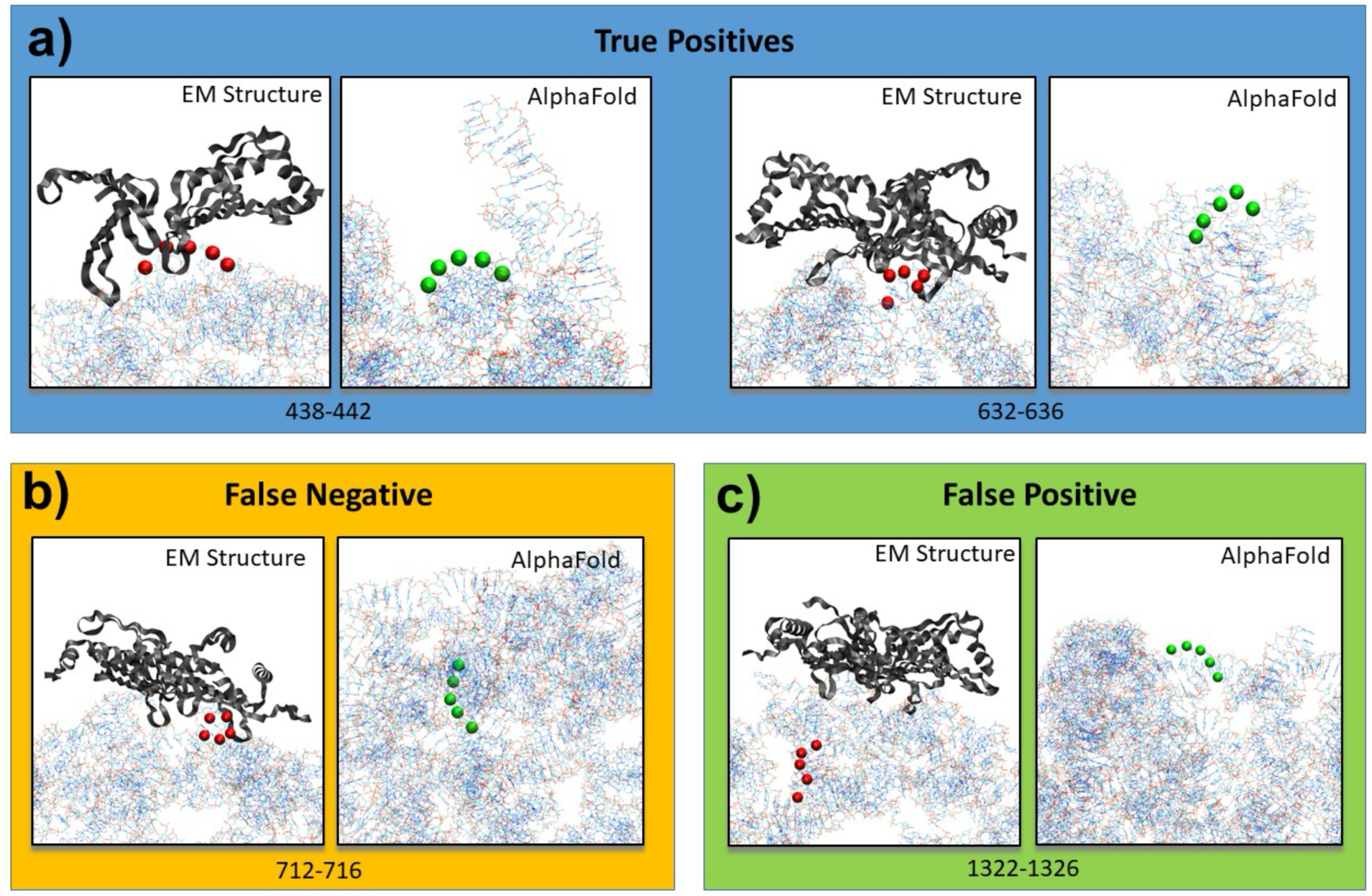
Representative examples of capsid protein binding site predictions using AlphaFold-predicted RNA structures. **(a)** Two true-positive examples in which experimentally determined capsid protein binding sites (red beads) were correctly predicted by the neural network. The corresponding RNA fragments (green beads) remain exposed in the AlphaFold-predicted structures, preserving local geometric features associated with capsid recognition. **(b)** A representative false-negative example. Although the RNA fragment is a capsid protein binding site in the cryo-EM structure, it is buried within the interior of the AlphaFold-predicted structure and is therefore incorrectly classified as a non-binding site. **(c)** A representative false-positive example. The RNA fragment does not contact capsid proteins in the experimentally determined structure but is exposed on the surface of the AlphaFold-predicted structure, resulting in an incorrect prediction as a binding site. These representative cases illustrate how differences between predicted and experimentally determined RNA conformations influence capsid protein binding site prediction.

Overall, these representative examples indicate that successful prediction of capsid protein binding sites requires not only static RNA structural information but also an understanding of the dynamic molecular processes that govern viral assembly.

## 4. Discussions

The selective recognition of viral RNA by capsid proteins is a fundamental step in the assembly of single-stranded RNA viruses. Identifying capsid protein binding sites is therefore essential for understanding the molecular mechanisms of viral genome packaging and has important applications in antiviral drug development, RNA therapeutics, and viral vector engineering. However, experimental identification of these binding sites remains labor-intensive and is available for only a limited number of viruses. In this work, we developed a computational framework that integrates RNA sequence, predicted tertiary structure, local geometric features, and machine learning to predict capsid protein binding sites directly from viral RNA sequences.

Using the Qβ bacteriophage as a proof-of-concept system, we demonstrated that local RNA geometry provides sufficient information for machine learning to identify capsid protein binding sites with good predictive performance. More importantly, our results showed that even approximate RNA local backbone structures predicted computationally preserve biologically meaningful local geometric information that can be exploited for functional prediction.

An important finding of this study is that predicted RNA tertiary structures retain measurable predictive information despite their limited structural accuracy. The moderate prediction performance obtained using AlphaFold-derived structures suggests that viral genome packaging is governed by both intrinsic RNA structural features and additional biological factors not captured by current computational models. While local RNA geometry contributes substantially to capsid recognition, static structural representations alone appear insufficient to fully explain binding specificity. RNA conformational dynamics, folding intermediates, structural ensembles, RNA modifications, molecular chaperones, and the physicochemical environment during viral assembly are all likely to influence RNA–protein interactions. These observations support a model in which local RNA geometry provides an important, but incomplete, encoding of the viral packaging code.

Despite the encouraging performance of the proposed framework, there remains considerable room for improvement. The current model represents each RNA fragment using five local geometric descriptors constructed solely from the phosphate (P) atoms of the RNA backbone. Although this coarse-grained representation efficiently captures local backbone geometry, it omits potentially important structural information associated with nucleotide bases, including base orientation, stacking, and accessibility, which may contribute to capsid protein recognition. Future models could therefore incorporate base-level conformational features together with backbone geometry to provide a more complete representation of local RNA structure. We can also adjust the length of RNA fragments. Moreover, additional information, including the nucleotide sequence, predicted secondary structure, and evolutionary conservation could be incorporated into the feature representation to provide a more comprehensive description of RNA–protein recognition. Furthermore, although a feedforward neural network achieved good predictive performance in this study, alternative machine learning algorithms, including support vector machines [20], random forests [21], or graph neural networks [22] may further improve prediction accuracy. Systematic comparison of these classifiers using the same benchmark dataset would provide valuable insights into the most effective computational strategy for this problem.

Another potential limitation of the present study lies in the construction of the benchmark dataset. Capsid protein binding sites were defined solely according to the minimum Euclidean distance between RNA nucleotides and capsid protein atoms in the cryo-EM structure with a fixed distance cutoff. While this criterion provides an objective and automated labeling strategy, a single short atom–atom contact may not necessarily represent a functionally meaningful RNA–protein interaction and may introduce nonspecific contacts into the benchmark dataset. A more physically informative definition could instead consider the buried or accessible surface area at the RNA– protein interface, which reflects the extent of molecular contact rather than relying on a single interatomic distance. Previous studies have shown that functional packaging signals are predominantly associated with stem-loop structures distributed throughout the viral RNA genome [12]. Future work should therefore incorporate biological knowledge, structural context, and experimental evidence to curate a more consistent benchmark dataset that better reflects functional packaging sites. Such improvements in label quality are expected to further enhance both the accuracy and the biological interpretability of machine learning models for capsid protein binding site prediction.

The present study focused exclusively on the Qβ bacteriophage as a proof-of-concept system. An important direction for future work is to extend the framework to additional single-stranded RNA viruses representing diverse viral families. As more experimentally resolved RNA–capsid complex structures become available, larger benchmark datasets can be constructed to train more robust and generalizable prediction models. Ultimately, we envision developing a general computational tool capable of predicting capsid protein binding sites directly from the genome sequences of diverse ssRNA viruses. Such a framework would facilitate studies of viral assembly, enable functional annotation of newly discovered viruses, and provide a computational platform for the rational design of antiviral strategies and RNA-based delivery systems.

## Acknowledgement

JZ is supported by the Center for Phage Technology, the NIH grant R01GM141659, TAMU ADM grant.

## Author Contributions

Y.M.W. and J.Z. designed research; Y.M.W. performed research; Y.M.W. analyzed data;

Y.M.W. and J.Z. wrote the paper.

